# Self Fourier shell correlation: properties and application to cryo-ET

**DOI:** 10.1101/2023.11.07.565363

**Authors:** Eric J. Verbeke, Marc Aurèle Gilles, Tamir Bendory, Amit Singer

## Abstract

The Fourier shell correlation (FSC) is a measure of the similarity between two signals computed over corresponding shells in the frequency domain and has broad applications in microscopy. In structural biology, the FSC is ubiquitous in methods for validation, resolution determination, and signal enhancement. Computing the FSC usually requires two independent measurements of the same underlying signal, which can be limiting for some applications. Here, we analyze and extend on an approach proposed by Koho et al. [1] to estimate the FSC from a single measurement. In particular, we derive the necessary conditions required to estimate the FSC from downsampled versions of a single noisy measurement. These conditions reveal additional corrections which we implement to increase the applicability of the method. We then illustrate two applications of our approach, first as an estimate of the global resolution from a single 3-D structure and second as a data-driven method for denoising tomographic reconstructions in electron cryo-tomography. These results provide general guidelines for computing the FSC from a single measurement and suggest new applications of the FSC in microscopy.

## 1 Introduction

The Fourier shell correlation (FSC) is defined as the normalized cross-correlation of corresponding shells between two signals in the frequency domain [2, 3]. In single particle electron cryo-microscopy (cryo-EM), the FSC has become the universal resolution metric and is used to assess the quality of a 3-D reconstruction [4, 5]. Additional major contributions of the FSC in cryo-EM include setting hyperparameters of iterative algorithms, as in 3-D refinement of structures [6], and estimation of the spectral signal-to-noise ratio (SSNR) [7, 8].

A core requirement of the FSC is the availability of two or more independent noisy measurements. In single particle cryo-EM, this is often achieved by splitting the data into random half sets [4]. However, for other forms of microscopy or data processing procedures, it is not always possible to apply the same strategy. To bypass the need for multiple measurements, a novel approach was recently proposed to estimate an FSC-like quantity from a single measurement, which we refer to as the self FSC (SFSC) [1]. This approach was initially used for image restoration in fluorescence microscopy [1] and has also been applied to estimating resolution in scanning electron microscopy [9]. The SFSC is implemented by first decimating an image in real space to produce downsampled images whose correlation with each other is then computed in Fourier space. While the interpretability of the original FSC has been discussed in [10, 11], the validity of the SFSC as a proxy for the FSC is more difficult to interpret, since the two downsampled signals are not independent.

In this work, we analyze the SFSC and give sufficient conditions on the statistics of both the signal and the noise under which the estimator is consistent with the standard FSC. Notably, we show that the assumptions required for the SFSC are more restrictive than the standard FSC and that use of the SFSC outside the defined conditions can give estimates that deviate significantly from the FSC. The conditions are easy to check and give practical guidelines to the applicability of the SFSC.

To demonstrate the validity of the SFSC, we provide two applications in the context of cryo-EM: first as a measure of the global resolution from a single map, and second as a data-driven method for denoising in electron cryo-tomography (cryo-ET). In the first application, we show that the resolution predicted by the SFSC from one half-map agrees with the standard FSC computed from two half maps, provided the conditions on the data described in this work are met. We then use the SFSC to denoise a reconstructed tomogram from cryo-ET data by applying a Wiener filter. Our approach provides significantly increased contrast and visibility compared to conventional low-pass filtering. The code used to generate the results in this work is available at: github.com/EricVerbeke/self_fourier_shell_correlation.

## 2 Results

Experimental evidence for the relationship between the correlation of noisy image measurements and the signal-to-noise ratio in electron microscopy date back to (at least) 1975 [12]. With the advent of the FSC, this relation was developed further to describe the decay in data quality with respect to spatial frequency based on a relation to the SSNR [7]. The SSNR is a central quantity in computational microscopy and has specific use in cryo-EM for denoising by Wiener filtering [13], and post-processing (*e*.*g*., 3-D structure sharpening) [4].

In this work, we consider the following simple model for estimating the FSC and therefore also the SSNR: we observe a single noisy measurement *y* of an underlying signal *x*:

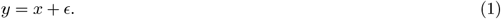

We discuss the effect of including the contrast transfer function (CTF) in the model in Appendix A.1. For the model in eq. (1), we assume that the ground truth signal is drawn from a mean-zero Gaussian distribution *x*∼ 𝒩 (0, Λ) with additive Gaussian colored noise *ϵ* ∼ 𝒩 (0, Σ), where Λ and Σ are the covariance matrices of the signal and noise respectively. We further assume that all entries of 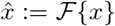 and 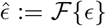, the discrete Fourier transforms (DFT) of *x* and *ϵ*, are statistically independent of each other, and thus have diagonal covariance matrices in the Fourier domain. That is, we have that 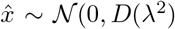 and 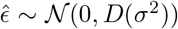, where *D*(*v*) ∈ ℂ^*d*×*d*^ denotes a diagonal matrix with entries *v* ∈ ℂ^*d*^, and *λ* and *σ* are vectors that are constant along frequency shells. The real space covariance matrices are therefore Λ = *F* ^∗^*D*(*λ*^2^)*F* and Σ = *F* ^∗^*D*(*σ*^2^)*F*, where *F* is the normalized DFT matrix and ^∗^ denotes the conjugate transpose. While the independence assumption on the signal in the Fourier domain may seem restrictive, it is typical in cryo-EM 3-D reconstruction—corresponding to a weighted *L*^2^ regularized problem in the maximum a posteriori formulation [6]; furthermore, it is justified in the “infinitely large” protein limit under the Wilson statistics model [14, 15]. The SSNR at spatial frequencies of radius *r* is then defined as:

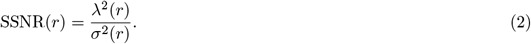

Typically, neither *λ*^2^ or *σ*^2^ are known *a priori* and thus must be estimated from data. In practice, the SSNR can be estimated if two independent and noisy measurements, *y*_1_ and *y*_2_, of the same signal *x* are available by computing their FSC.

The FSC is defined as:

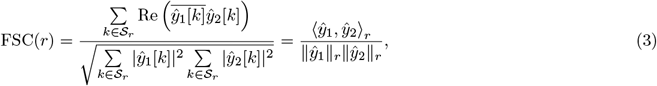

where *ŷ*_1_ and *ŷ*_2_ are the DFT of *y*_1_ and *y*_2_, the overline denotes the complex conjugate, and, ⟨·, ·⟩_*r*_ denotes the standard inner product on ℂ^*d*^ restricted to the shell 𝒮 _*r*_ with ∥ · ∥_*r*_ being the associated norm. We use brackets to denote indexing of a discrete function and *k* for the multi-index on a Fourier grid. The link between the FSC and SSNR is made by considering a related deterministic quantity, denoted EFSC:

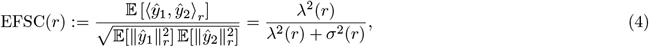

where 𝔼 is the expectation. We note that while 𝔼 [FSC(*r*)] ≠ EFSC(*r*), the estimated quantity has proven to be a useful proxy for the SSNR. From the EFSC, we see that the SSNR can be estimated as:

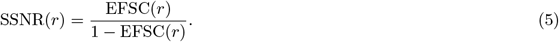

It is a common practice to replace the EFSC by the empirical FSC eq. (3) computed from two signals to estimate the SSNR.

### 2.1 Fourier shell correlation from a single measurement

The computation of the standard FSC requires two independent measurements of a signal. In this work, the goal is to estimate the FSC and SSNR from a single measurement. A solution proposed in [1] is to compute the FSC from downsampled versions of the same measurement. This approach is originally implemented by first taking a noisy, real-space measurement and decimating into a checkerboard-like pattern to form half-sized approximations of the original measurement, as shown in Figure A2. The FSC between pairs of downsampled signals can then be computed.

Here, we modify the downsampling procedure such that the real-space measurements are split into even and odd terms along one spatial dimension at a time, thus providing two downsampled versions for each dimension. The FSC is then computed between each downsampled measurement pair for each dimension and the reported FSC is taken to be the average, as shown in Figure 1. There are two main advantages of this approach compared to the checkerboard-like splitting pattern. First, our scheme preserves the Nyquist frequency for each axis except the one split into even and odd terms. Second, we show in Appendix A.2 that splitting in a checkerboard-like pattern scales the variance of the noise in the SFSC by 2^dim^ where dim is the number of axes split into even and odd terms. For the 1-D case, a measured signal is decimated by simply splitting into even and odd terms.

**Figure 1.**
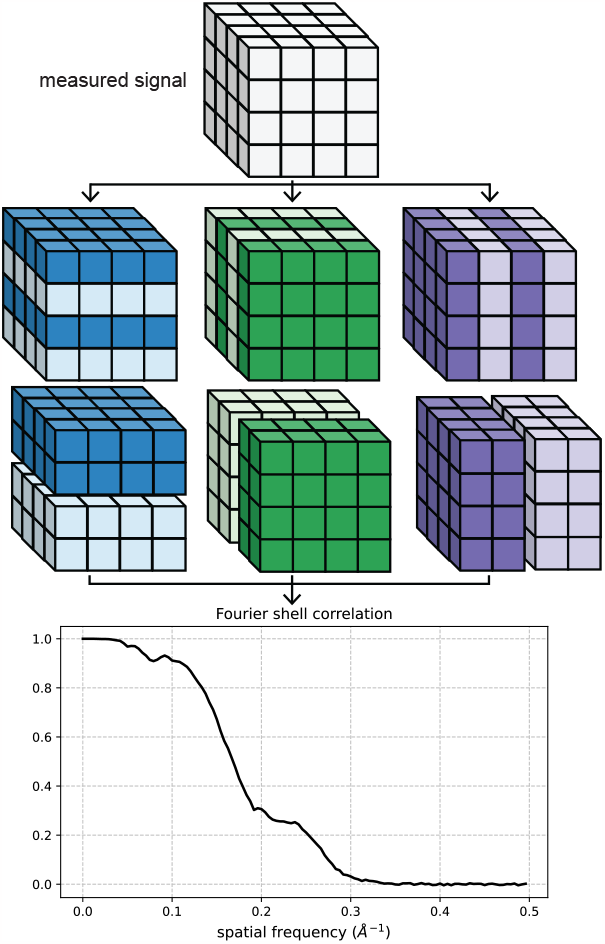
Illustration of the Fourier shell correlation computed from a downsampled signal (*i*.*e*., the SFSC). The measured signal is split into even and odd voxels for each dimension and the SFSC is computed between the respective pairs. The reported FSC is taken to be the average of the three pairs.

While this process is always easily computable, it is not clear that the estimate is meaningful. Indeed, the basis of the connection between the FSC and SSNR is the statistical independence of two measurements. However, in the SFSC case, the two measurements are simply downsampled versions of the same noisy measurement which are correlated in any practical scenario. Despite the apparent correlation, we show that under conditions on the statistics of both the signal and the noise, the SFSC may still be used to estimate the SSNR from the downsampled measurement, which can be used to infer the SSNR of the original measurement.

### 2.2 Conditions for accurate estimation of the FSC from the SFSC

We present our main analysis for the SFSC here using the one-dimensional case for simplicity, although we show in Appendix A.3 that it naturally extends to higher dimensions. Following the model in eq. (1), let *y* be a discrete 1-D measurement of length *N*, where we assume *N* is even. The measurment *y* is then downsampled by splitting it into even index terms *y*_*e*_[*n*] = *y*[2*n*] and odd index terms *y*_*o*_[*n*] = *y*[2*n* + 1] for *n* ∈ {0, …, (*N/*2) − 1}. The DFT of the even and odd term measurements can be related to the DFT of the original measurement *y* as follows (see Appendix A.3 for derivation):

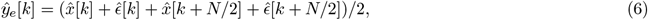

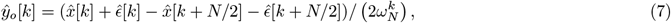

where *ŷ*_*e*_ and *ŷ*_*o*_ are the DFTs of *y*_*e*_ and *y*_*o*_, and *ω*_*N*_ = exp(−2*πi/N*). We note that if the higher frequency terms are small (*i*.*e*., there is a rapid decay in the power spectrum), then *ŷ*_*e*_[*k*] and *ŷ*_*o*_[*k*] are approximately equal after a phase shift of *ŷ*_*o*_[*k*] by 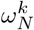. Thus, as noted in [1], when computing the SFSC between downsampled pairs, a phase shift correction must be included. That is:

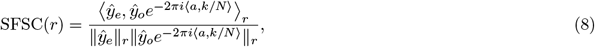

where *a* denotes the translation. We discuss the origin and effect of this translation further in Appendix A.4.

Our goal is to show that the SFSC is approximately equal to the FSC such that it can also provide an estimate of the SSNR as in eq. (5). Following the same arguments as stated for the EFSC in eq. (4), we have that:

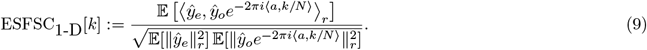

Under the standard assumption that the signal and the noise are statistically independent, we then get:

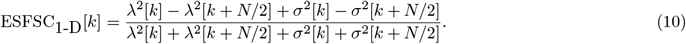

Thus, in general, the estimates for the EFSC and ESFSC are not the same. However, if we consider the following two assumptions:

#### Assumption 1

The Gaussian noise distribution is white, namely *σ*^2^[*k*] = *σ*^2^[0] ∀*k*,

#### Assumption 2

The power spectrum of the signal decays such that *λ*^2^[*k*] ≫ *λ*^2^[*k* + *N/*2]
then, we have that:

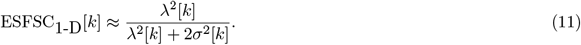

That is, the ESFSC approximates the EFSC with an additional doubling on the variance of the noise. Given the above assumptions are met, the ESFSC can be related to the EFSC by:

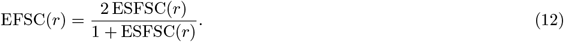

We illustrate the importance of the assumptions on the signal and noise in Figure 2, and how to circumvent these assumptions in Figure 3 using a synthetic 2-D image as an example. The clean image originates from a projection of the 3-D structure of a human gamma-aminobutyric acid receptor (available as entry EMD-11657 in the electron microscopy data bank) [16]. Using a typical B-factor decay in cryo-EM [4], we generate each image as 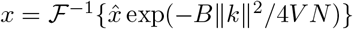, where *V* is the voxel (or pixel) size and *B* modulates the decay. Here the image size is *N* ×*N* = 360× 360 with a pixel size of 0.81Å. Noisy measurements are then produced by adding Gaussian noise. To generate noise which decays with spatial frequency (*i*.*e*., such that Assumption 1 is broken), a B-factor decay may also be applied to the additive noise and is delineated as *B*_*signal*_ and *B*_*noise*_ when necessary.

**Figure 2:**
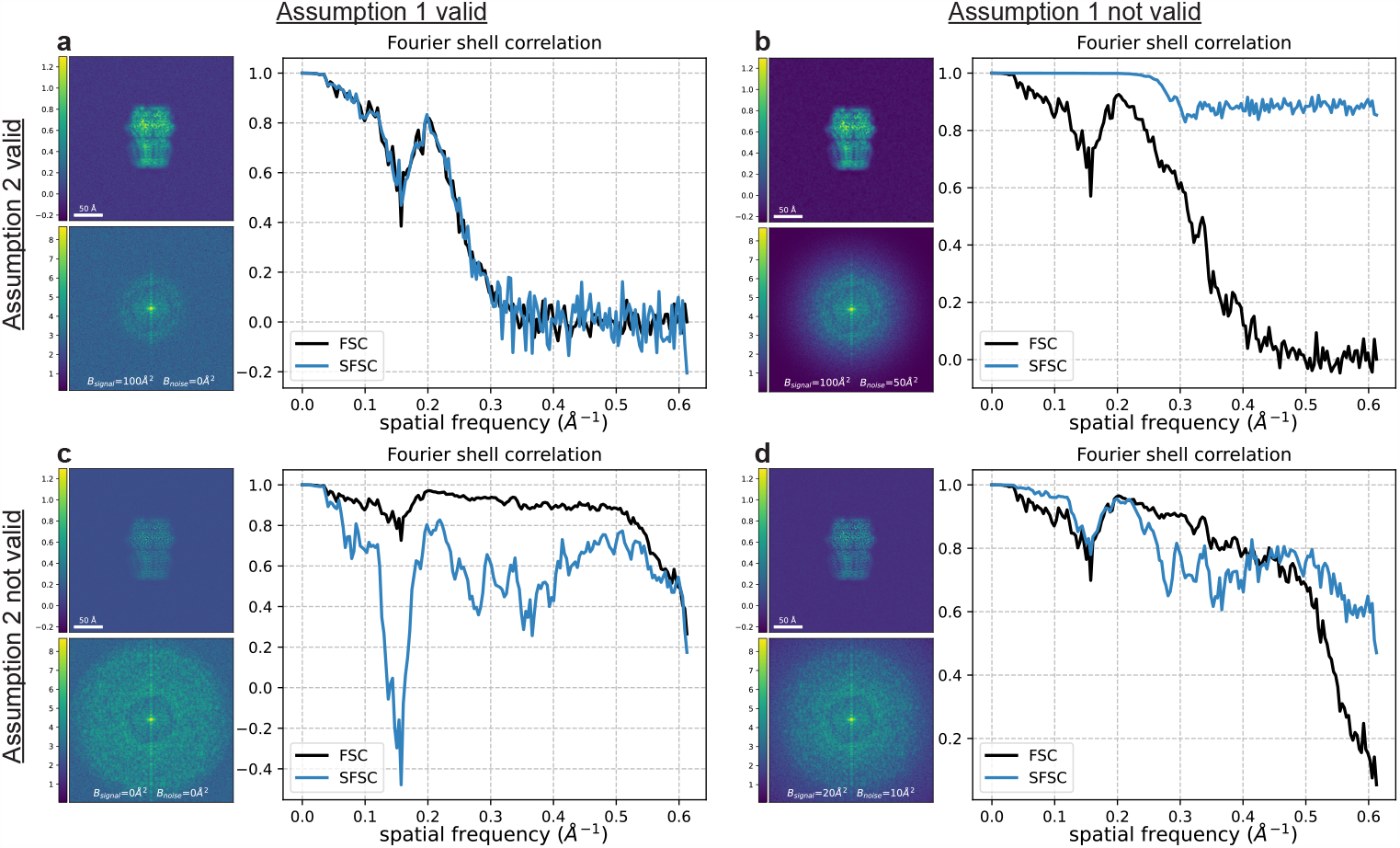
Conditions on the statistics of the signal and noise under which the SFSC accurately estimates the FSC. Each panel shows the image, power spectrum and associated SFSC for a signal that satisfies or fails to satisfy both Assumption 1 and Assumption 2. The SNR was set to 15 for each image with additive Gaussian noise that decays with spatial frequency when specified. The FSC was computed for each case using two synthetic images generated with the same parameters but independent noise. **(a)** *B*_*signal*_ = 100Å ^2^, *B*_*noise*_ = 0 Å ^2^. Both assumptions are met and the SFSC accurately estimates the FSC. **(b)** *B*_*signal*_ = 100Å ^2^, *B*_*noise*_ = 50 Å ^2^. The noise is not white Gaussian and the SFSC overestimates the FSC. **(c)** *B*_*signal*_ = 0 Å ^2^, *B*_*signal*_ = 0 Å ^2^. The noise is white Gaussian but the signal does not have rapid decay. The SFSC underestimates the FSC. **(d)** *B*_*signal*_ = 20 Å ^2^, *B*_*noise*_ = 10 Å ^2^. Neither assumption is met and the SFSC fails to estimate the FSC. This figure demonstrates that the naive SFSC provides an accurate estimate of the FSC only if Assumption 1 and Assumption 2 are met.

**Figure 3:**
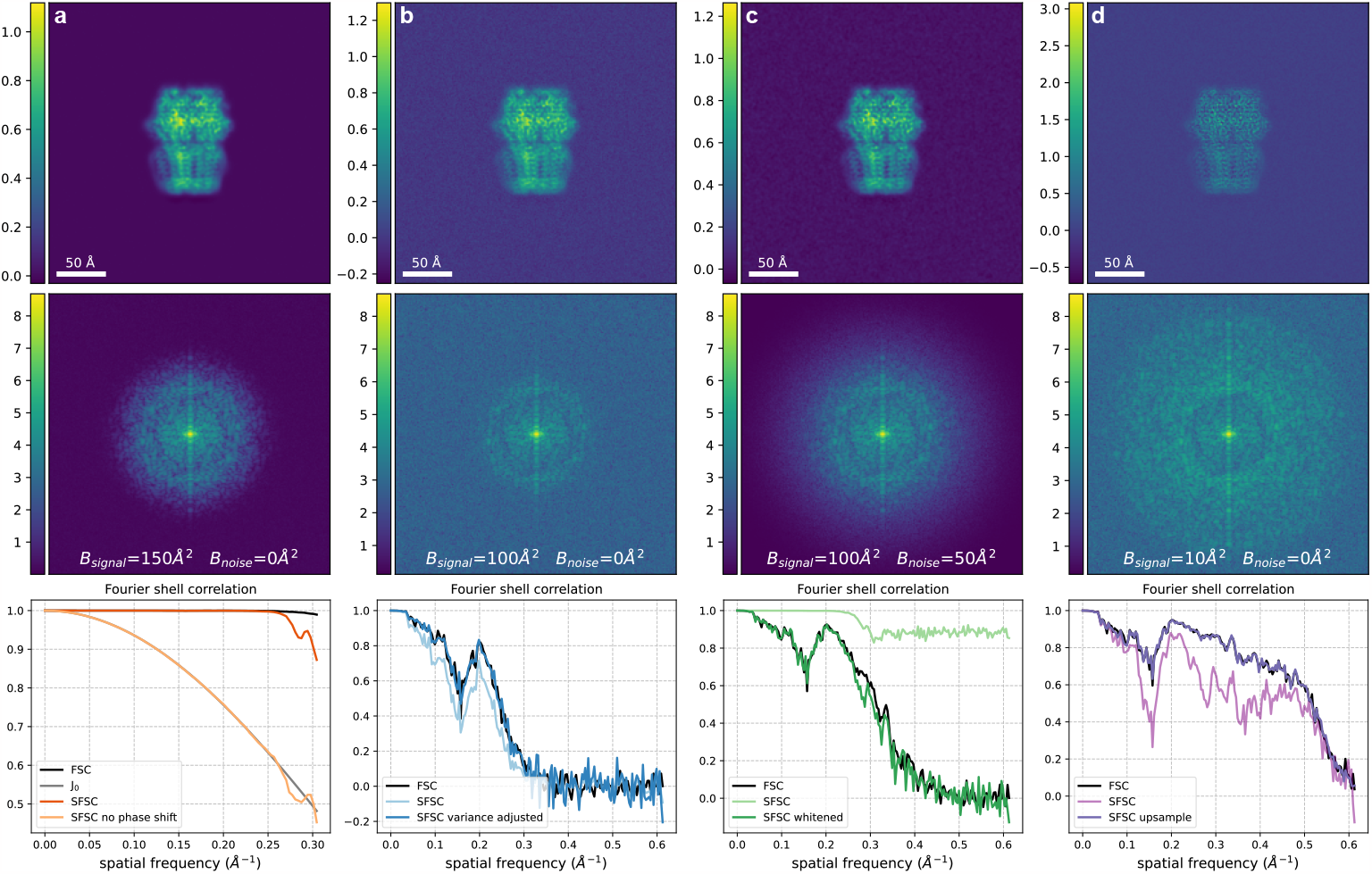
Corrections required for the SFSC to accurately estimate the FSC. The image and corresponding power spectrum in each column were generated with a specified SNR and a B-factor on both the signal and noise to exemplify each case. The FSC was computed for each case using two synthetic images generated with the same parameters but independent noise. **(a)** Phase shift correction, SNR = 10^5^, *B*_*signal*_ = 150Å ^2^, *B*_*signal*_ = 0Å ^2^. If the phase shift induced by downsampling is not corrected, the 2-D SFSC reduces to *J*_0_, a scaled zeroth order Bessel function of the first kind (see Appendix A.4). **(b)** Correction for the scaled variance, SNR = 15, *B*_*signal*_ = 100Å ^2^, *B*_*noise*_ = 0Å ^2^. Both assumptions on the signal and noise are met. The SFSC estimates the FSC according to eq. (12) after adjusting for the scaled variance. **(c)** Whitening transform, SNR = 15, *B*_*signal*_ = 100Å ^2^, *B*_*noise*_ = 50Å ^2^. After applying a whitening transform, the SFSC estimates the FSC. **(d)** Upsampling, SNR = 10, *B*_*signal*_ = 10Å ^2^, *B*_*noise*_ = 0Å ^2^. If the signal does not have rapid decay but has been whitened, the SFSC estimates the FSC only after upsampling. These correcting factors extend the applicability of the SFSC.

Given both assumptions on the data are met, and that the correction for the scaling of the variance in eq. (12) is applied, we show in Figure 2a that the SFSC accurately estimates the FSC. If the noise is not white Gaussian, but instead decays with spatial frequency, the SFSC is unreliable and overestimates the FSC (Figure 2b). However if the noise is white Gaussian, but the power spectrum of the signal does not have rapid decay, then the SFSC underestimates the FSC (Figure 2c). Finally, if neither assumption is met, the SFSC fails to approximate the FSC and tends to give an overestimate (Figure 2d). These results underpin the behavior of the SFSC, whether or not it should be applied, and motivate the improvements to the algorithm described in the following sections which circumvent the assumptions.

### 2.3 Accounting for colored noise in the SFSC

Microscopy images are often contaminated by colored Gaussian noise. Specifically, in cryo-EM, noise is often modeled by a covariance matrix that is diagonal in the Fourier domain, but with entries that vary [17]. In this case, Assumption 1 is violated. From eq. (10), the ESFSC approximates the EFSC only in the case of white Gaussian noise and should not be expected to match otherwise. However, in the scenario where the noise is not white but its distribution can be estimated, we can first *whiten* the measurement prior to computing the SFSC. Suppose that the Fourier transform of the noise distribution is 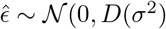). We define the Fourier transform of the whitened noisy measurement *ŷ*_*w*_ as:

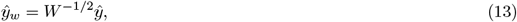

where *W* = *D*(*σ*^2^). By construction, the noise in *ŷ*_*w*_ is white:

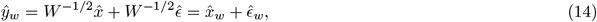

where 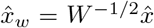 and 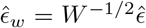, and 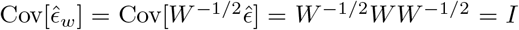. While this transform changes the signal, the SSNR of the new signal is the same as the original one. Noise whitening of data has statistical justifications which are described in [18].

To demonstrate the effect of colored noise on the SFSC, we generated an image with a noise spectrum that decays following exp(− *B* ∥ *k* ∥^2^ */*4*V N*), with *B* = 50Å^2^. We show in Figure 3c that after applying a whitening transform, we can recover the FSC from the SFSC. While this procedure can always be done if the noise level can be estimated, we note that it also leads to a simpler scheme for estimating the SSNR. Specifically, if the noise variance can be estimated, the ratio of the noise variance subtracted from the power spectrum to the noise variance is, in expectation, also equal to the SSNR. In fact, this is always possible and yields approximately equivalent results to the standard FSC. The noise level is typically computed as part of the cryo-EM reconstruction process and could be used for a more direct measure of the SSNR without the need of the FSC. We demonstrate this simpler approach for estimating the SSNR in Appendix A.5.

### 2.4 SFSC for measurements with slow decaying spectrum

Assumption 2 requires that there is rapid decay in the power spectrum of the underlying signal. If this is not the case, we can introduce an additional correction to the SFSC. We propose the following approach: upsample the measurement by zero-padding in Fourier space to increase the length of the measurement to *Ñ* = 2*N*, then subtract off the noise level from the numerator. The effect of the zero-padding is to set the high frequency terms to zero, and thus their variance is also zero (*i*.*e*., *λ*^2^[*k* + *Ñ /*2] = *σ*^2^[*k* + *Ñ /*2] := 0). Returning to eq. (9) we see:

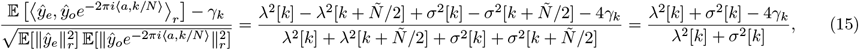

where *γ*_*k*_ is a value we have chosen. The above equation equals the desired EFSC in eq. (4) when we set 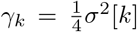. Importantly, this procedure only works after a whitening transform of the original measurement since the variance of the noise is known. We show in Figure 3d that upsampling a whitened measurement with a slow decaying power spectrum recovers the expected correlation curve. The effect of upsampling, and more generally frequency filtering prior to computing the SFSC, is discussed in Appendix A.6.

### 2.5 Estimating resolution from a single cryo-EM map

For a conventional 3-D reconstruction pipeline in single particle cryo-EM, the data are split into random half sets to generate two independent half maps which are used to compute the FSC. To verify the assumptions and corrections introduced in this work, we show that the global resolution can be estimated from a single cryo-EM half map using the SFSC. We use the 3-D structures of a 20S proteasome (EMD-24822 [19]), a 70S ribosome (EMD-13234 [20]) and two small membrane proteins (EMD-27648 [21], EMD-20278 [22]) as examples.

After 3-D reconstruction, we do not expect the noise to be white, but instead to increase proportionally to ∥*ξ*∥ due to the Fourier slice theorem, whereas the signal will show a strong exponential decay due to the B-factor. Thus, after whitening, our method applies. However, deposited maps usually have masking in either the volume or images which impact the noise statistics. Here we use the the noise in the corners of the reconstructions to estimate the noise distribution, but this assumes no masking was used. Otherwise, the noise in the corners and center will display different statistics. Specifically, we use the region outside a sphere which encompasses the molecular structure to estimate the noise by computing the spherically averaged power spectrum, defined as 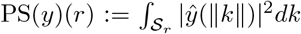 (see Figure A4). We then apply the whitening transform and upsample procedure before computing the SFSC. The resolution reported using the standard FSC and the resolution calculated from the SFSC at a threshold of 1*/*7 [4] are approximately equal (Figure 4a-c) except for the case where there is a non-unifom noise distribution (Figure 4d). We attribute the deviation of the SFSC from one at the low frequencies to the difficulty in accurately estimating the noise using *ad hoc* methods. These results suggest that the SFSC provides a viable alternative for estimating resolution in cryo-EM and could be used in the absence of half maps.

**Figure 4:**
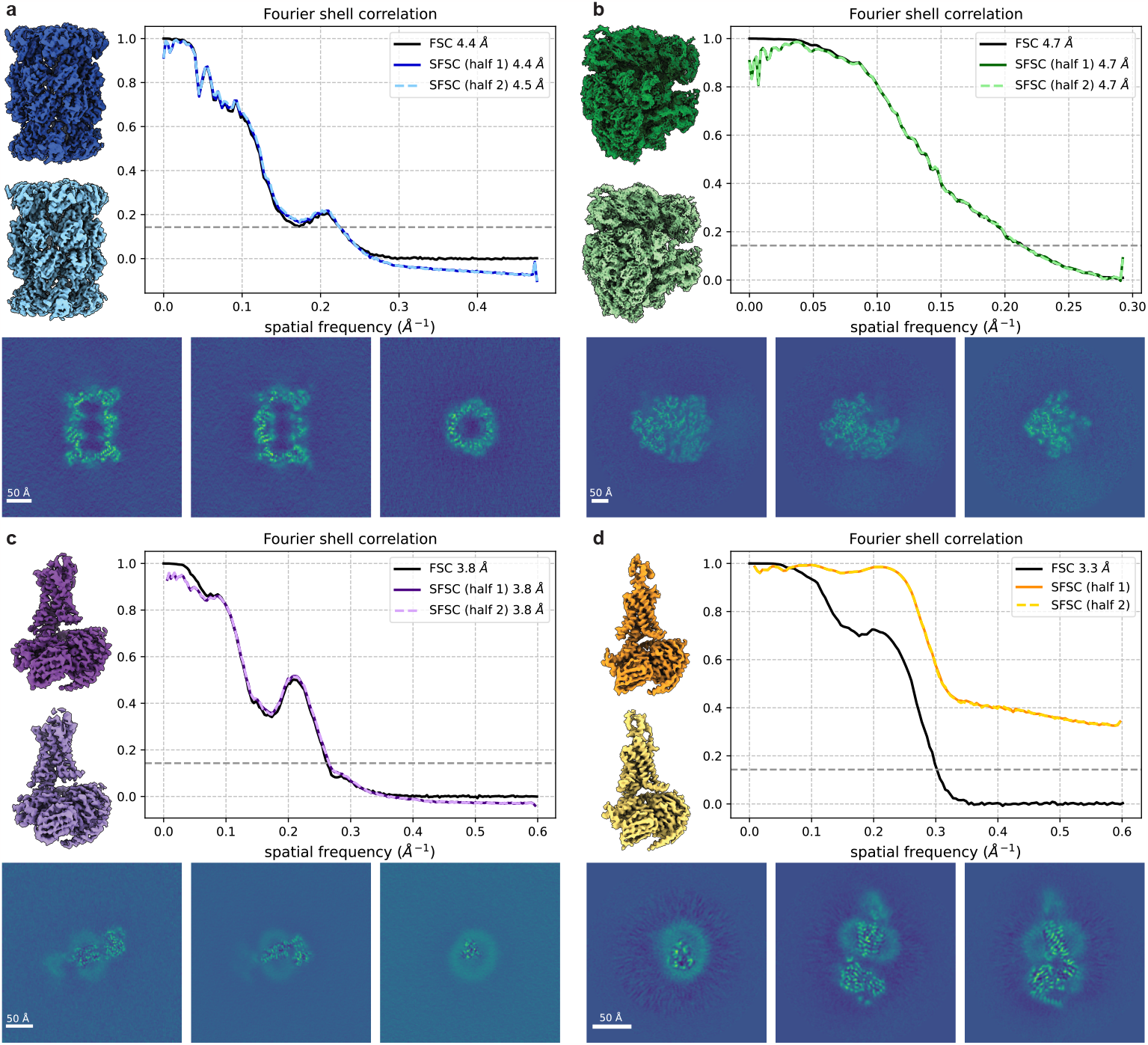
Global resolution estimates from single maps. The SFSC is computed for each half map after applying the noise whitening and upsampling procedure. The noise is estimated by computing the spherically averaged power spectrum from the region outside a sphere encompassing the structure. The SFSC is approximately equal to the standard FSC for **(a)** EMD-24822 (grid points = 360^3^, voxel size = 1.05Å), **(b)** EMD-13234 (grid points = 336^3^, voxel size = 1.7Å) and **(c)** EMD-27648 (grid points = 416^3^, voxel size = 0.83Å), but fails for **(d)** EMD-20278 (grid points = 288^3^, voxel size = 0.83Å) due to the non-uniform noise which can be seen in the central slice images.

### 2.6 Denoising a reconstructed tomogram

Having established the necessary assumptions and corrections under which the SFSC provides an estimate of the SSNR, we next demonstrate an application to denoising tomographic reconstructions from cryo-ET data. In a typical cryo-ET tilt series data collection scheme, projection images are measured at ± 60° in several degree increments. The recorded frames at each tilt are then motion corrected, aligned and a reconstruction technique is used to generate the 3-D tomogram. Due to the low electron beam dose required for imaging biological samples, the SSNR in cryo-ET data is low [23], and so there is a need for denoising methods [24]. Additionally, unlike in single particle cryo-EM, there are no related measurements to boost the SSNR by averaging. Thus, cryo-ET provides an ideal use case for the SFSC. Alternative approaches for estimating the resolution in cryo-ET such as computing the FSC from reconstructions of the even and odd images in a tilt series are described in [25].

Considering the model for a noisy measurement in eq. (1), the minimum mean square error estimator for *x* given *y* under the Gaussian assumptions is known as the Wiener filter, and is widely used in cryo-EM [13, 26, 27]. The Wiener filter is defined as:

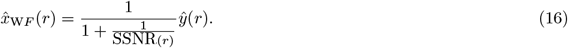

In cryo-ET, a common practice is to provide an *ad hoc* SSNR for Wiener filtering, or simply to use a low-pass filter. However, given the relationship between the EFSC and SSNR in eq. (5), and that the SFSC can estimate the FSC, we show that the SFSC provides a simple, data-driven method for applying a Wiener filter. Combining eq. (16) with eq. (5), we get that 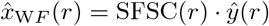. Effectively, each shell in Fourier space for measurement *y* is weighted according to the correlation profile from the SFSC. The idea of self-Wiener filtering has already been suggested in the context of signal processing and is described in [28].

Here we show in Figure 5 that applying a Wiener filter from the computed SFSC improves the visibility of a reconstructed tomogram. The data in this example is *C. elegans* tissue from EMD-4869 [29]. In order to accurately estimate the SSNR, we know from Assumption 1 that the noise must be white Gaussian. To estimate the noise variance for a subsection of the tomogram, we select a slice above the region of interest. We then compute the SFSC and apply the Wiener filter in eq. (16). The resulting denoised section of the tomogram shows enhanced contrast over the original and a low-pass filtered version. Specifically, the ribosomes and membrane edges stand out from the background. We additionally consider CTF effects and compare our approach to the noise learning method cryo-CARE [30] using a different data set (EMD-15056 [31]) in Appendix A.8. These results demonstrate that the SFSC can provide a simple, data-driven and parameter-free filter for improving the visualization of tomograms.

**Figure 5:**
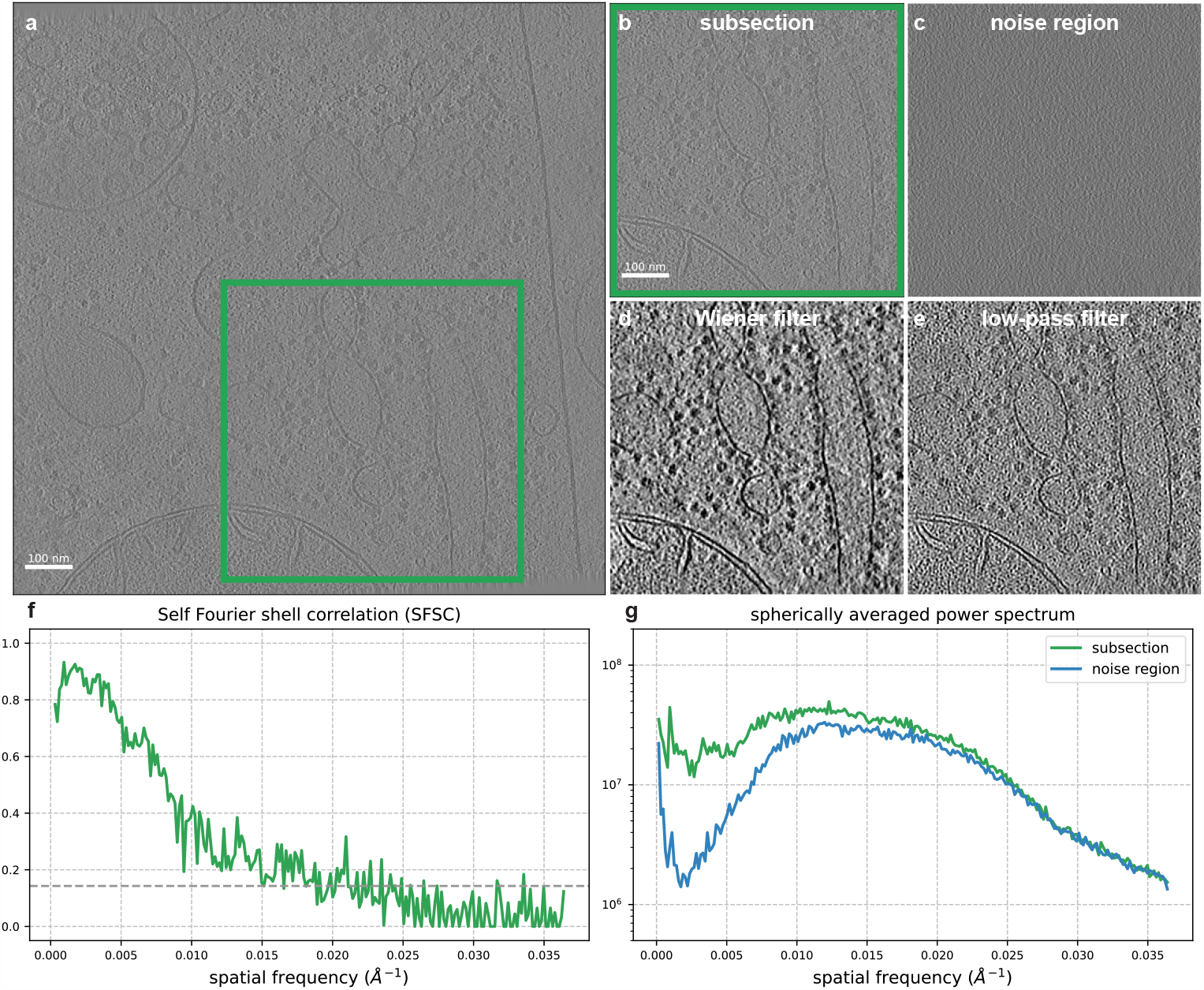
Denoising a reconstructed tomogram using the SFSC. **(a)** Slice of a reconstructed tomogram of *C. elegans* tissue from EMD-4869 (*N* ×*N* = 928 × 928, pixel size = 13.7Å). **(b)** Region of interest from a subsection of the tomogram (*N* ×*N* = 464 ×464). **(c)** Slice of the tomogram selected vertically above the region of interest containing background noise. **(d)** Slice from the region of interest after applying a Wiener filter. **(e)** Conventional low-pass filter of the subsection at 66Å determined using the 1*/*7 threshold of the SFSC. Both the Wiener filtered and low-pass filtered images are displayed at a threshold of ± 2 standard deviations of the pixel values. **(f)** SFSC computed from the tomogram subsection. **(g)** Spherically averaged power spectrum of the region of interest slice and the background noise slice. The Wiener filter computed from the SFSC provides significantly increased contrast compared to a low-pass filtering approach.

## 3 Discussion

In this work, we derive a set of conditions and corrections required for accurately computing the Fourier shell correlation from downsampled versions of a single noisy measurement. We demonstrate that we are able to estimate the global resolution from a single map and denoise a reconstructed tomogram using the SFSC. Our approach is broadly applicable and allows for estimation of the SSNR if it is not possible to collect replicate measurements or use prior information. Furthermore, our approach does not require instrument specific calibration as described in [1]. The corrections we introduce in this work extend the applicability of the SFSC but also suggest a simpler path to estimating the SSNR provided an estimate of the noise can be obtained. We show that the same logic applies for any data processing pipeline in cryo-EM that estimates the noise or computes half maps (see Appendix A.5). If the noise cannot be accurately estimated or is non-uniform, then the SFSC should not be expected to work. While we estimate the noise with *ad hoc* methods here, using more accurate approaches typically employed in cryo-EM data processing pipelines could improve the SFSC and associated Wiener filter.

Although the SFSC is not always applicable, there are many situations that can benefit from having an estimate of the SSNR from a single measurement. For example, in single particle cryo-EM, there are a growing number of methods which generate 3-D structures from manifold embeddings and do not produce independent half maps with which to compute the standard FSC [32–34]. Thus there is a need for alternative methods to estimate signal and noise statistics. In principle, the SFSC could also be used to circumvent splitting data into half sets during 3-D reconstruction. This has the potential to lead to improved reconstructions due to an increase in SSNR from using the full data set. One additional application of the SFSC to single particle cryo-EM could be to both denoise and estimate the resolution of 2-D class averages.

Other useful applications of the SFSC could include validation of SSNR enhancement after modification by neural network based approaches [35]. Similarly, the SFSC could provide an alternate measurement of the SSNR for noise learning based methods [30, 36, 37]. While the noise learning approaches give impressive results for denoising tomograms (see Figure A6), the benefit of the Wiener filter presented here is that it is fast to compute, requires no parameter tuning, and does not require extra storage (*e*.*g*., from reconstructing tomograms using odd and even frames). Further analysis of the SFSC for use with cryo-ET could also account for the missing wedge as well as directional and local resolution effects.

## 4 Methods

### 4.1 Main algorithm

The algorithm presented in this work consists of 3 main steps and 3 preprocessing steps, depending on the properties of the measured signal. The main steps are parameter-free and can be written succinctly as: 1) for each dimension, split the measured signal into even and odd index terms along that dimension, 2) for each pair of downsampled measurements, compute the SFSC, and 3) average the SFSC from all pairs.

### 4.2 Data preprocessing

Prior to computing the main algorithm, the user should first discern if Assumption 1 and Assumption 2 are met. This can be checked by plotting the spherically averaged power spectrum. If both assumptions are met, the power spectrum at the latter half of spatial frequencies should appear approximately constant. However, if the required assumptions are not met, the preprocessing steps described in this work should be applied. These steps can be applied regardless of the signal and noise properties as long as an estimate of the noise variance can be obtained. The preprocessing steps are: 1) estimate the the noise variance, 2) whiten the measured signal, 3) upsample the whitened signal.

### 4.3 Estimating the noise variance

Several strategies exist to estimate the noise variance from data. This estimate is required for computing the SFSC if the noise is not white Gaussian. For the case of estimating the noise variance from a half map of a 3-D reconstruction in cryo-EM, we use an *ad hoc* approach by taking the region outside a sphere encompassing the molecular structure. For example, with EMD-24822, we use a spherical mask with radius = 115Å. The noise variance is then estimated by computing the spherically averaged power spectrum of the noise region.

### 4.4 Denoising

To estimate the noise variance for a reconstructed 3-D tomogram, we use a slice of the tomogram above the region of interest. Although the noise slice does not reflect the true 3-D noise profile, and more accurate methods can be used, we show that it is a suitable estimate for whitening based on the results of Wiener filtering. When applying the Wiener filter, we find that an additional low-pass filter at the spatial frequency corresponding to the 1*/*7 threshold from the SFSC can subtly enhance the contrast.

## Data Availability

All data sets used in this work are available from the Electron Microscopy Data Bank [38]. The entries used are EMD-11657, EMD-24822, EMD-13234, EMD-27648 and EMD-20278 for molecular structures, and EMD-4869 and EMD-15056 for reconstructed tomograms.

## Code Availability

The source code used to produce the results and figures in this work is available at github.com/EricVerbeke/self_fourier_ shell_correlation. The SFSC will be made available as a tool in the software package ASPIRE [39].

## Acknowledgements

E.J.V., M.A.G. and A.S. are supported in part by AFOSR under Grant FA9550-20-1-0266, in part by Simons Foundation Math+X Investigator Award, in part by NSF under Grant DMS-2009753, and in part by NIH/NIGMS under Grant R01GM136780-01. T.B. is partially supported by the NSF-BSF award 2019752, the BSF grant no. 2020159, and the ISF grant no. 1924/21. We thank Zunlong Ke and Ricardo D. Righetto for valuable discussion and insight regarding cryo-ET.

## Author contributions

E.J.V. and A.S. conceived of this project. E.J.V. developed the software and ran numerical experiments. E.J.V., M.A.G., T.B. and A.S. designed the experiments and analyzed the results. E.J.V. and M.A.G. wrote the manuscript. All authors commented on and edited the manuscript.

## A Appendix

### A.1 Applicability of the SFSC to CTF modified signals

Microscopy images are typically corrupted by imaging artifacts. In cryo-EM, this is modeled by the contrast transfer function (CTF) which is the Fourier transform of the point spread function of the microscope. We therefore discuss the effects of these modifications on the estimation of the FSC by SFSC. Often, the forward model may be written as:

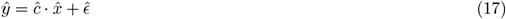

where 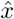 is the Fourier transform of the underlying signal, *ĉ* is the CTF effect, and *ŷ* is the Fourier transform of the corrupted observation. Given two images of the same signal affected by the same CTF, the FSC can be readily used to estimate the SSNR, but it should be noted that it will be an estimate of the SSNR for the corrupted signal 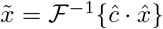, and not the clean signal *x*. Similarly, the SFSC can also be used to estimate the SSNR of the corrupted signal assuming the measurement follows the properties assumption 1 and assumption 2. We illustrate this with two simulated examples. The first example is for a typical CTF in cryo-EM images, described by:

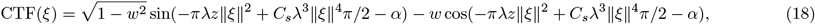

with *w* = 0.1, *λ* = 2.51 pm, *z* = 2.8 *μ*m, *C*_*s*_ = 2.0 *μ*m, and *α* = 0.87. The second example is for a CTF that resembles a tilt series image in cryo-ET (see Appendix A.8), described by:

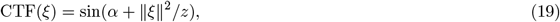

with *α* = 3*/*4 and *z* = 5. We generate noisy, CTF corrupted images following eq. (17). The FSC, SFSC and spherically averaged power spectrum are then computed for each image Figure A1. As expected, the oscillations of the CTF modulate the correlation profile in both the FSC and SFSC. Importantly, the SFSC still approximates the FSC well. In these scenarios, the traditional resolution value obtained from a threshold may not be meaningful as we expect the FSC to oscillate. Most importantly, while the CTF may change the reported resolution value, the Wiener filter computed using the SFSC of a CTF modified signal is still a statistically optimal filter for denoising.

We note that if the CTF is not radially symmetric, then the CTF corrupted signal should not be expected to satisfy radially symmetric assumptions. In this case, both the FSC and SFSC are poor estimators of the SSNR. However, when applied to reconstructions from a collection of CTF corrupted images with a random and uniform distribution of poses, the reconstructions will have approximately radially symmetric noise and variance even if the images do not.

**Figure A1:**
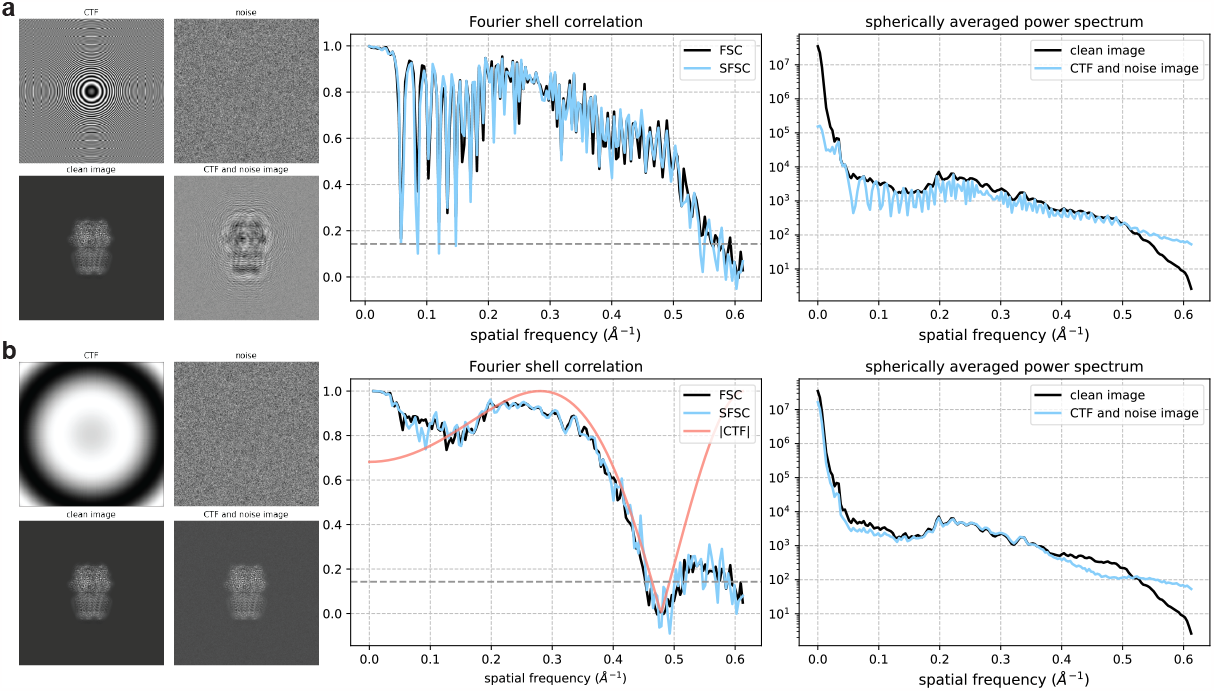
Effect of the CTF on the FSC and SFSC. Clean image is a projection of EMD-11657 (*N* × *N* = 360 × 360, pixel size = 0.812Å). Additive Gaussian noise was generated with SNR = 15 and *B*_*noise*_ = 10 Å ^2^. **(a)** Results for image formed using CTF in eq. (18). **(b)** Results for image formed using CTF in eq. (19). Also plotted is the absolute value of the radial CTF.

### A.2 Comparison of downsampling methods

The foundation of the SFSC is that a real space measurement can be downsampled by decimation to generate multiple approximations of the measurement whose correlation can then be computed in Fourier space. In our proposed downsampling scheme (see Figure 1), the number of measurements to be compared is equal to the spatial dimensions of the signal (*e*.*g*., 3 pairs for a 3-D signal). In the procedure proposed by Koho et al. [1], there are 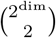 pairs to be compared. There are two main disadvantages of splitting in a checkerboard-like pattern compared to only splitting along one dimension at a time, as proposed in this work. First, the variance of the noise in the downsampled measurements is scaled by 2^dim^, where dim is the number of dimensions split across. This relation is derived in the following section. Second, the Nyquist frequency will be reduced to half of the original. We demonstrate both of these effects using the 2-D case of an image in Figure A2.

**Figure A2:**
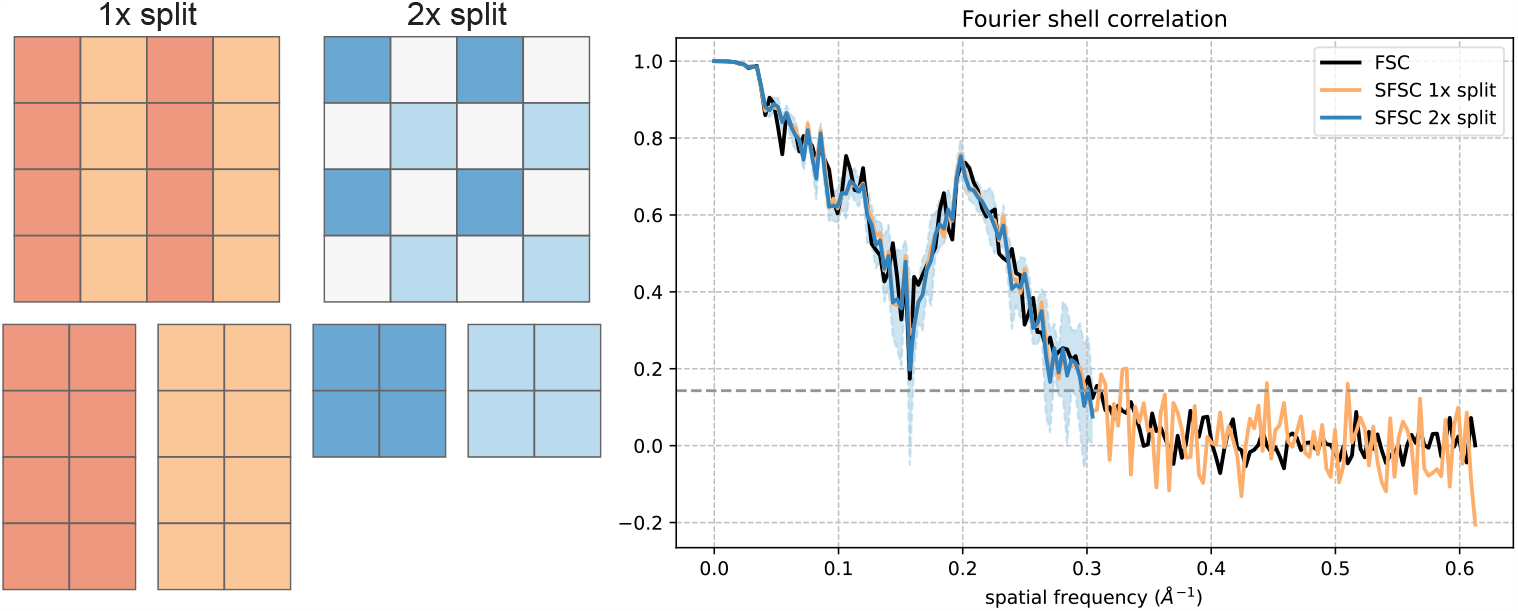
SFSC computed from an image downsampled by splitting along 1 and 2 dimensions. The pixel diagram depicts one pair out of two for the 1 × split and one pair out of six for the 2 × split. The input image used for the FSC and SFSC is from Figure 3B. The SFSC from the 1 × split image is reported as the average of both pairs. The SFSC from the 2 × split image is reported as the average of the six combinations and is plotted with an error envelope representing the standard deviation of the SFSC from all combinations.

### A.3 General relation between FSC and SFSC

To relate the SFSC to the FSC, we need to relate the DFT of the decimated signals to the DFT of the original signal. We first show this relation for a signal split along one dimension, as advocated in this work, and then provide an example for a signal split along two dimensions, as done for images in the original splitting scheme by Koho et al. [1], to demonstrate how it extends to a signal split along multiple dimensions.

Let *x* be a 1-D signal of length *N*. The DFT of *x* is defined as 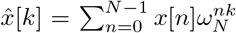, where *ω*_*N*_ = exp(−2*πi/N*). The signal *x* can be split into even index terms *x*_*e*_[*n*] = *x*[2*n*] and odd index terms *x*_*o*_[*n*] = *x*[2*n* + 1] for *n* ∈ {0, …, (*N/*2) − 1}. We want to relate 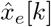 and 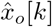, the DFTs of *x*_*e*_ and *x*_*o*_, to the DFT of the original signal 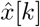. The relation can be seen by splitting the DFT of *x* into the sum of the even and odd terms:

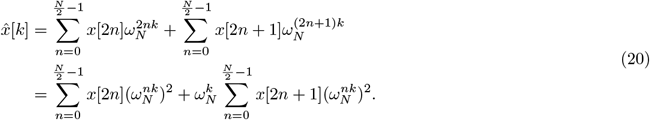

Next, we use that 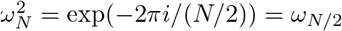 to get:

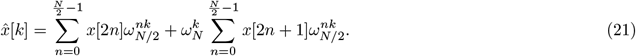

Applying the definition of the DFT to the right side of the equation yields:

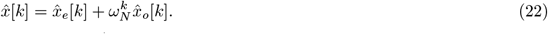

Importantly, *k* = 0, …, *N* − 1, and 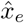 and 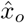 are *N/*2 periodic^1^. We can then independently relate 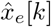 and 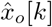 to 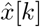 as follows:

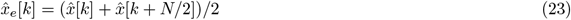

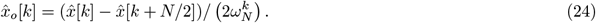

Equation (23) and eq. (24) form the framework of our analysis presented in Section 2.2, and describe the relation between the DFTs of a 2-D or 3-D measurement that has been split into alternating voxels along one dimension.

The analysis above holds for a signal split along multiple dimensions as well, since the DFT can be applied along subsequent dimensions. Specifically, we are referring here to the checkerboard-like splitting pattern from [1]. Suppose now that *x* is a 2-D signal (*i*.*e*., an image). If the signal *x* is split into even and odd index terms along both dimensions, then we have that:

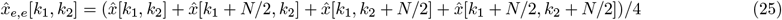

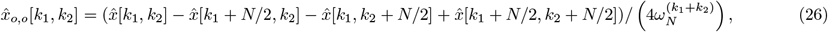

where *e* denotes even and *o* denotes odd indexing for each dimension of *x*, and *k*_1_ and *k*_2_ are the frequency indexes of 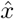. In the case of a noisy measurement, we are interested in the relation between the SFSC from the downsampled measurements using this splitting scheme and the SSNR. Following the arguments presented in Section 2.2, if both assumptions on the signal and noise are met, then:

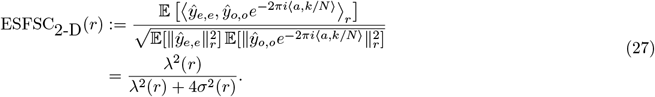

For the general case of decimating into even and odd terms over multiple dimensions, we see that:

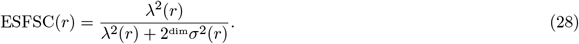

Thus, when splitting along multiple dimensions, there is a scaling of 2^dim^ on the noise variance compared to a scaling of 2 from splitting once as proposed in this work. From eq. (28), the relation between the EFSC and ESFSC split over multiple dimensions is:

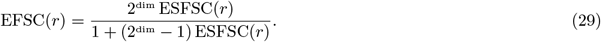

We show in Figure 3A that the SFSC will yield an underestimate of the FSC if the correction in eq. (29) is not applied. For the 2-D case in Figure 3A, the SFSC was computed using our splitting scheme. Thus eq. (29) is equal to eq. (12).

**Figure A3:**
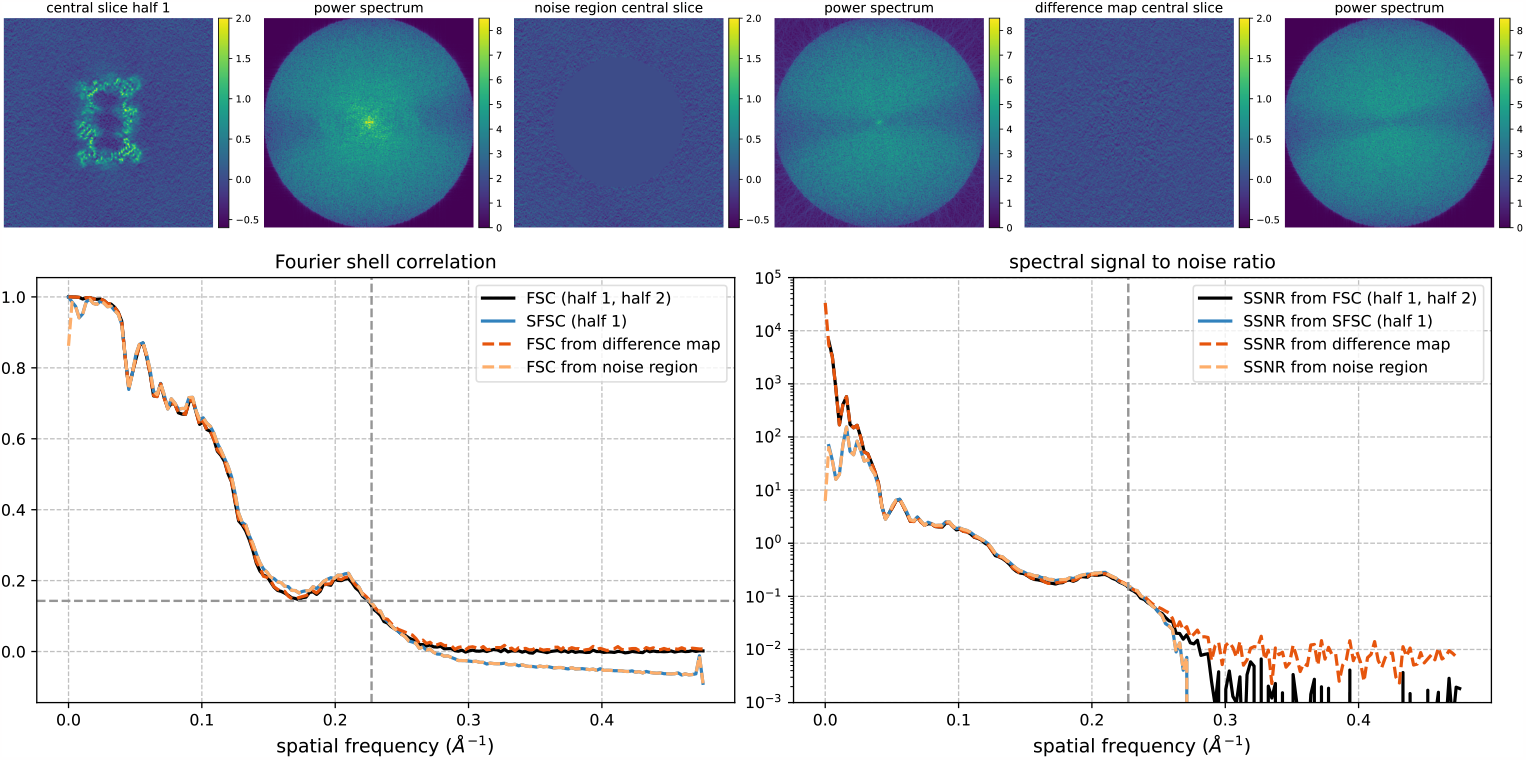
Comparison of the FSC from half maps and the SSNR estimated from measurement noise. The FSC and SSNR are computed from EMD-24822 by four different methods and show similar profiles.

### A.4 FSC with a phase shifted input

We consider the effect of a translation between real space signals when computing the FSC. A translation is naturally induced between pairs of decimated measurements when computing the SFSC and must be corrected. For example, as noted in [1], when an image is decimated in a checkerboard-like pattern, each downsampled image pair is offset by a single pixel in each dimension. Here we show that if the power spectrum of the signal is approximately spherically symmetric, a translation between the inputs to the SFSC, and more generally the FSC, leads to a signal-independent quantity unique for the 1-D, 2-D and 3-D case.

Continuing with the 2-D case, the decimated images represent the same area as the original image, but with half the length for each split dimension. This leads to an effective pixel size of twice the original. With respect to the original image, we expect the translation between the downsampled image pairs to be:

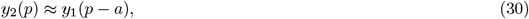

where *a* = [1*/*2, 0]^*T*^ for an image split along one dimension, or *a* = [1*/*2, 1*/*2]^*T*^ for an image split along two. A translation of 1*/*2 reflects the increase in pixel size. If the power spectrum of the image decays fast, then the adjacent pixels are indeed correlated. Here we consider the signal to be continuous, with its Fourier transform defined as *ŷ*(*ξ*) = *y*(*p*) exp(−2*πi ξ, p*)*dp*, where *ξ* is the frequency variable. A translation in real space is equivalent to a phase shift in Fourier space. It follows that:

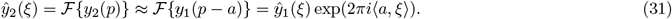

That is, we expect the spectrum of image *y*_2_ to be approximately equal to the spectrum of image *y*_1_ multiplied by a phase shift. The continuous analogue of the FSC can be expressed as:

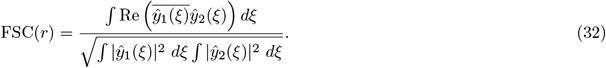

From eq. (30), *ŷ*_1_ and *ŷ*_2_ are approximately equal after a phase shift. We can then rewrite the FSC as:

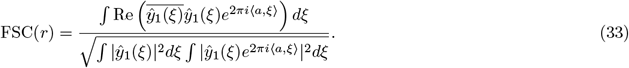

If the power spectrum of *y* is approximately spherically symmetric, then we then have that:

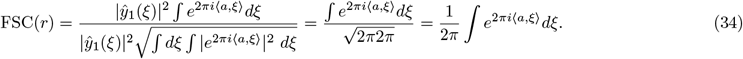

For the 2-D case, the inner product can be written as: ⟨*a, ξ*⟩ = ∥*a*∥ ∥*ξ*∥ cos(*ϕ*) = ∥*a*∥ *r* cos(*ϕ*). Since the integral is over the ring, we can reparametrize *ϕ*, the angle between *a* and *ξ*, so that cos(*ϕ*) = sin(*θ*). We then get that:

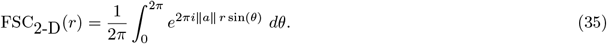

In this form, the FSC is equivalent to a scaled zeroth order Bessel function of the first kind, *J*_0_(*q*), defined as:

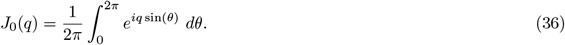

Comparing eq. (35) to eq. (36), we see that FSC_2-D_ (*r*) = *J*_0_(2*π* ∥*a*∥ *r*). We show in Figure 3A that failing to account for the induced phase shift in a 2-D image reduces the SFSC to eq. (36) which no longer matches the FSC. For the 3-D case, under the similar assumption of a spherically symmetric power spectrum, evaluating the integral in eq. (34) yields:

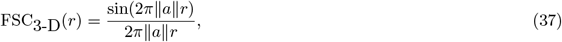

which is equivalent to a scaled and normalized sinc function. Finally, in the 1-D case, without accounting for the translation, we simply have that the FSC reduces to exp(−2*πik/N*). Apart from describing a necessary correction needed for computing the SFSC, these results emphasize that the inputs to the FSC must be carefully aligned. Otherwise, the output from the FSC might not reflect the signal to noise ratio, but rather the deterministic and signal-independent functions described above.

### A.5 Estimation of the SSNR without the FSC

In Section 2.2 we describe the assumptions under which the SFSC provides an estimate of the FSC. For some scenarios, if both Assumption 1 and Assumption 2 hold, we can estimate *σ*^2^, the variance of the noise, directly from the high frequencies of the power spectrum. In particular, this is true for high frequency shells when the signal spectrum has decayed and the noise level dominates the power spectrum (*i*.*e*., the SSNR is small). If the spherically averaged power spectrum does not appear flat at high frequencies, the noise variance cannot be estimated from the power spectrum and a different approach is needed. Estimating the noise is a standard part of the cryo-EM pipeline and is essential for the whitening transform described in Section 2.3. In this work we show the noise variance can be estimated from regions of noise or the difference of half maps from 3-D reconstructions. If one has access to the noise variance, the following simpler and more direct approach can be used to obtain the SSNR:

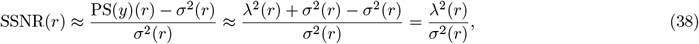

from which the EFSC can also be computed as:

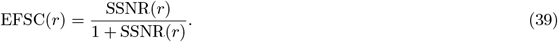

We show in Figure A3 that the SFSC from a half map and the FSC determined using a noise region of the half map for eq. (38) are approximately equal. We additionally show that if two half maps have been computed, the SSNR, and therefore the FSC, can be estimated just as well from the difference of half maps. Thus, this property is not intrinsic to just the SFSC. Nonetheless, the FSC still has the advantage of being scale invariant and can be applied even when there is ambiguity in the scale between the two measurements.

### A.6 Remarks on frequency filtering for the SFSC

To avoid aliasing when downsampling a signal, the standard approach is to first low-pass filter the original signal [40]. However, frequency filtering should not be applied when computing the SFSC as both the low and high frequency terms of the original signal are needed to correctly estimate the FSC. This can be directly seen in the DFTs of the downsampled signals which are modified by 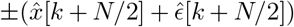, the higher frequency terms. If an ideal low-pass filter was applied to the original signal such that all frequencies *k > N/*4 are set to zero, the resulting SFSC would equal 1 at all frequencies, regardless of the SSNR. Similarly, if the original signal was high-pass filtered such that all frequencies *k < N/*4 are set to zero, the SFSC would equal − 1 at all frequencies. Thus, in order to accurately estimate the FSC from the SFSC, frequency filtering of the original signal should be avoided. The upsampling procedure in Section 2.4 used to compute the SFSC for measurements without decaying power spectrum effectively creates a low-pass filter. However, since the measurement must first be whitened for the upsampling procedure to work, we can compensate for the necessary high frequency noise terms since the variance of the noise has been set to 1.

### A.7 Noise estimation from reconstructed volumes

**Figure A4:**
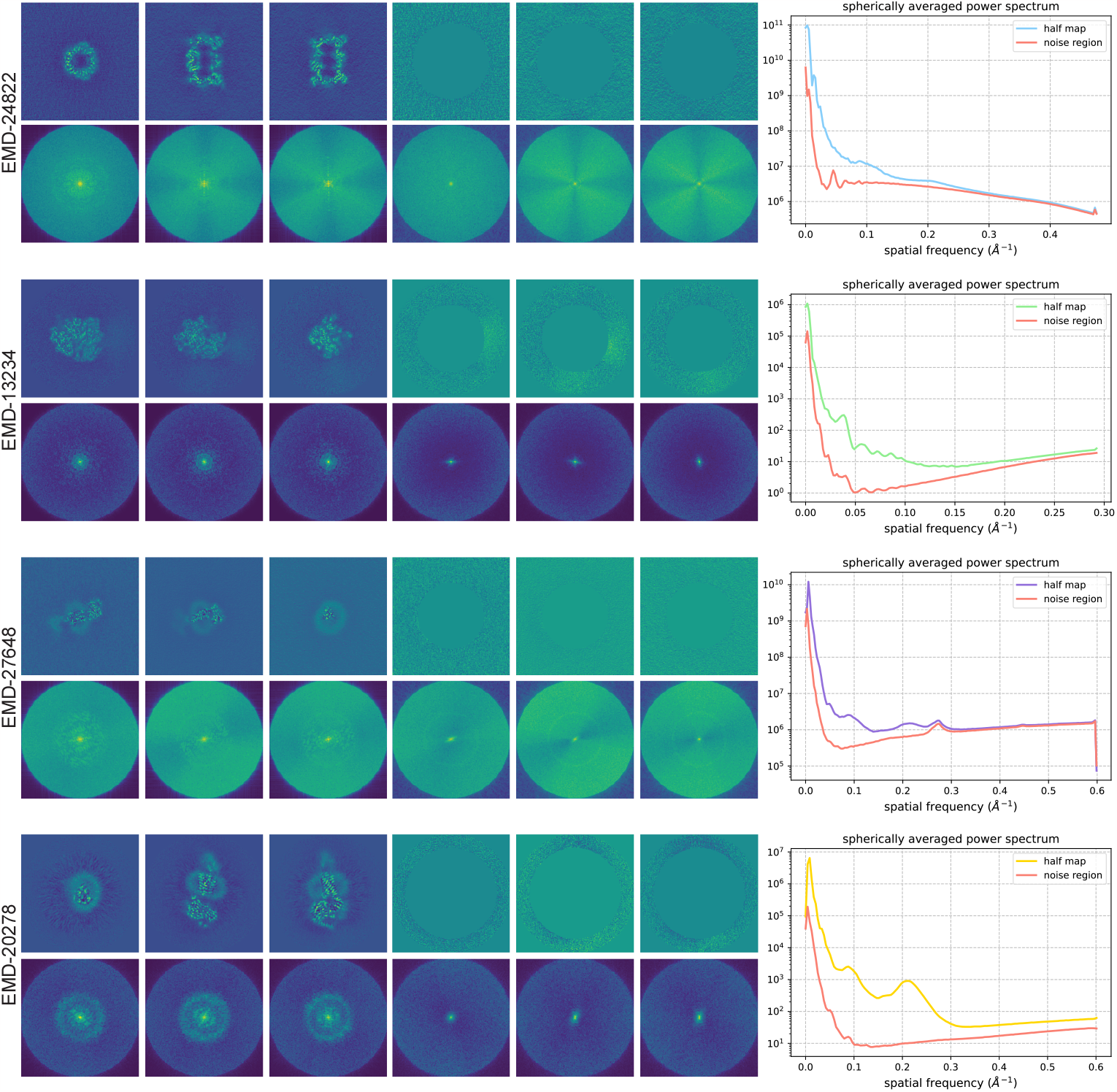
Central slices of the raw half maps, the regions containing only noise, and their corresponding power spectrum (below each image) for the four structures in Figure 4.

**Figure A5:**
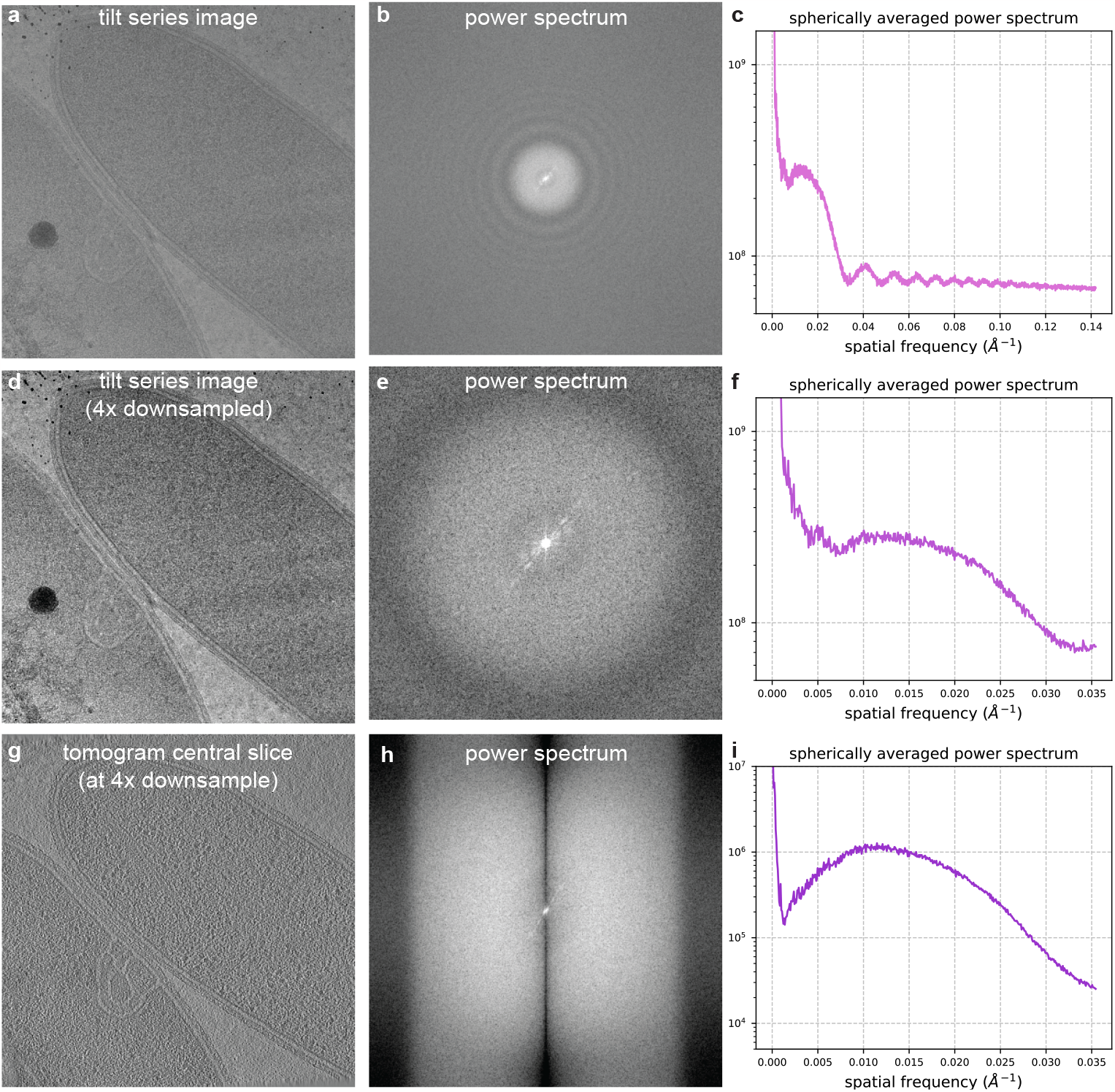
CTF visualization for tilt series and reconstruction. **(a)** Motion corrected tilt series image from EMPIAR-11058 (*N × N* = 3712 *×* 3712, pixel size = 3.52Å). **(b)** Power spectrum of tilt series image. **(c)** Spherically averaged power spectrum of tilt series image. **(d)** Tilt series image in **(a)** downsampled 4*×* as done in the reconstruction pipeline. **(e)** Power spectrum of the downsampled tilt series image. **(f)** Spherically averaged power spectrum of the downsampled tilt series image. **(g)** Central slice of the reconstructed tomogram. **(h)** Power spectrum of a central slice of the reconstructed tomogram. **(i)** Spherically averaged power spectrum of the reconstructed tomogram central slice. The tilt series image and downsampled version used for reconstruction clearly show the CTF. However the CTF effects are less visible in the reconstructed tomogram.

**Figure A6:**
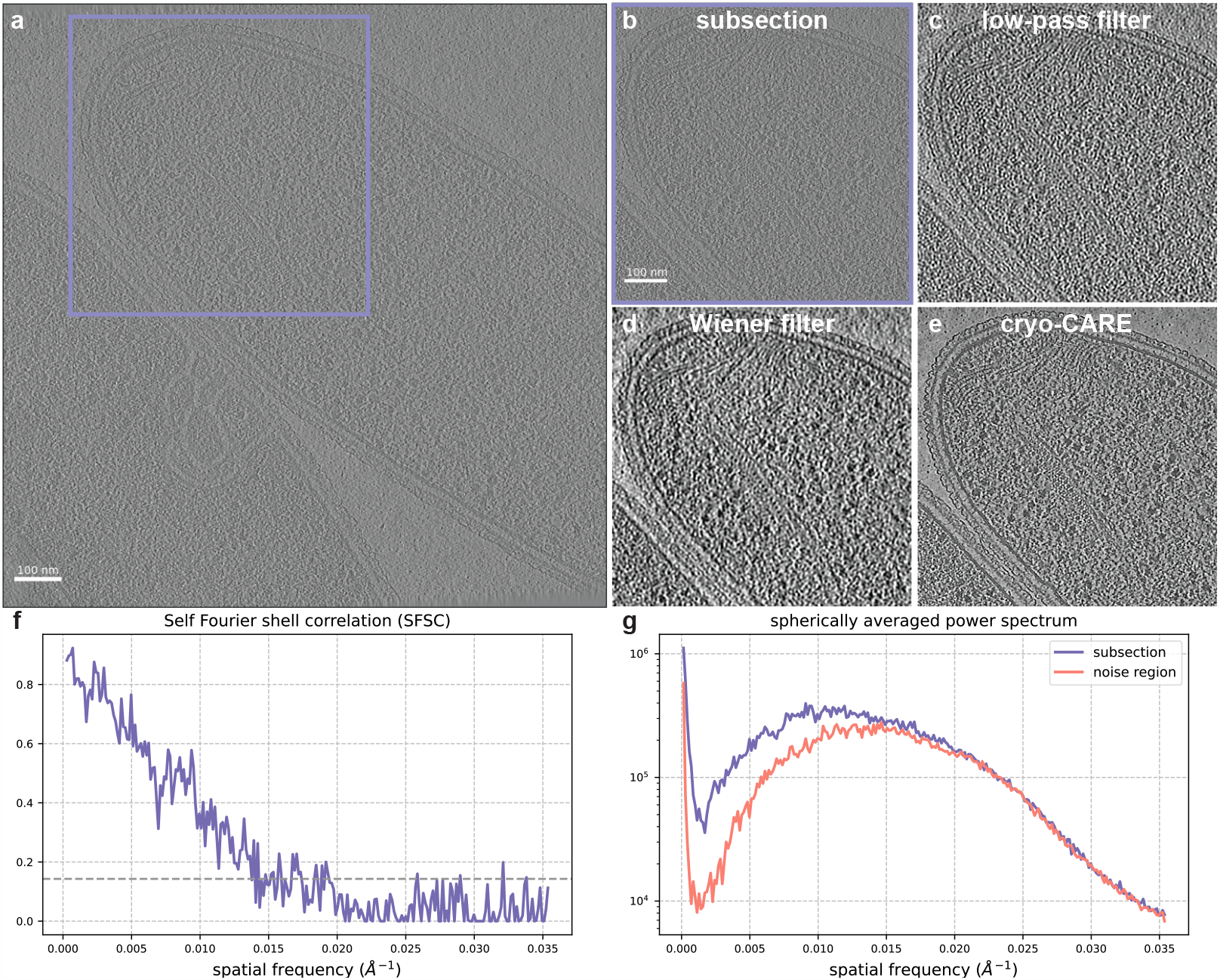
Comparison of denoising methods for a subsection of the tomogram EMD-15056 [31]. **(a)** Slice of the reconstructed tomogram (*N × N* = 928 × 928, pixel size = 14.1Å). **(b)** Region of interest from a subsection of the tomogram (*N× N* = 500*×* 500). **(c)** Low-pass filter of the subsection at 72Å, corresponding to the 1*/*7 threshold in the SFSC. **(d)** Slice from the region of interest after applying a Wiener filter. **(e)** Slice from the region of interest denoised using cryo-CARE. Images in (c-e) are displayed at a threshold of ± 2 standard deviations of the pixel values. **(f)** SFSC computed from the tomogram subsection. **(g)** Spherically averaged power spectrum of the region of interest slice and the background noise slice. While the Wiener filter improves contrast for features like membrane edges and ribosomes, cryo-CARE excels at both suppressing background and enhancing relevant high frequency information.

### A.8 CTF and denoising for tomograms

In Figure A5, we visualize the CTF from a tilt series image and the tomographic reconstruction of EMD-15056 [31]. The power spectrum of the full size tilt series image clearly displays Thon rings. However, the downsampled image used for reconstruction only shows one zero crossing of the CTF. The Wiener filter denoised tomogram and a comparison to the cryo-CARE denoised tomogram is shown in Figure A6.

## Footnotes

1 This well known recursive identity is at the core of the fast Fourier transform algorithm.

## References

1. Koho, S. et al. Fourier ring correlation simplifies image restoration in fluorescence microscopy. Nature Communications 10, 3103 (Dec. 2019).

2. Harauz, G. & Van Heel, M. Exact filters for general geometry three dimensional reconstruction. Optik 73, 146–156 (Feb. 1986).

3. Saxton, W. O. & Baumeister, W. The correlation averaging of a regularly arranged bacterial cell envelope protein. Journal of Microscopy 127, 127–138 (Aug. 1982).

4. Rosenthal, P. B. & Henderson, R. Optimal Determination of Particle Orientation, Absolute Hand, and Contrast Loss in Single-particle Electron Cryomicroscopy. Journal of Molecular Biology 333, 721–745 (Oct. 2003).

5. Scheres, S. H. W. & Chen, S. Prevention of overfitting in cryo-EM structure determination. Nature Methods 9, 853–854 (Sept. 2012).

6. Scheres, S. H. RELION: Implementation of a Bayesian approach to cryo-EM structure determination. Journal of Structural Biology 180, 519–530 (Dec. 2012).

7. Unser, M., Trus, B. L. & Steven, A. C. A new resolution criterion based on spectral signal-to-noise ratios. Ultramicroscopy 23, 39–51 (Jan. 1987).

8. Penczek, P. A. Three-dimensional spectral signal-to-noise ratio for a class of reconstruction algorithms. Journal of Structural Biology 138, 34–46 (Apr. 2002).

9. Perdigão, L. M. A. et al. Okapi-EM: A napari plugin for processing and analyzing cryogenic serial focused ion beam/scanning electron microscopy images. Biological Imaging 3, e9 (2023).

10. van Heel, M. & Schatz, M. Fourier shell correlation threshold criteria. Journal of Structural Biology 151, 250–262 (Sept. 2005).

11. van Heel, M. & Schatz, M. Reassessing the Revolution’s Resolutions. bioRxiv (Nov. 2017).

12. Frank, J. & Al-Ali, L. Signal-to-noise ratio of electron micrographs obtained by cross correlation. Nature 256, 376–379 (July 1975).

13. Penczek, P. A. in Methods in Enzymology 35–72 (Elsevier, 2010).

14. Gilles, M. A. & Singer, A. A molecular prior distribution for Bayesian inference based on Wilson statistics. Computer Methods and Programs in Biomedicine 221, 106830 (June 2022).

15. Wilson, A. J. C. Determination of absolute from relative X-Ray intensity data. Nature 150, 152–152 (Aug. 1942).

16. Nakane, T. et al. Single-particle cryo-EM at atomic resolution. Nature 587, 152–156 (Nov. 2020).

17. Bendory, T., Bartesaghi, A. & Singer, A. Single-particle cryo-electron microscopy: Mathematical theory, computational challenges, and opportunities. IEEE signal processing magazine 37, 58–76 (2020).

18. Leeb, W. & Romanov, E. Optimal spectral shrinkage and PCA with heteroscedastic noise. IEEE Transactions on Information Theory 67, 3009–3037 (May 2021).

19. Sae-Lee, W. et al. The protein organization of a red blood cell. Cell Reports 40, 111103 (July 2022).

20. Xue, L. et al. Visualizing translation dynamics at atomic detail inside a bacterial cell. Nature 611, E13–E13 (Nov. 2022).

21. Faust, B. et al. Autoantibody mimicry of hormone action at the thyrotropin receptor. Nature (Aug. 2022).

22. Liang, Y.-L. et al. Toward a Structural Understanding of Class B GPCR Peptide Binding and Activation. Molecular Cell 77, 656–668.e5 (Feb. 2020).

23. Koster, A. J. & Bárcena, M. in Electron Tomography: Methods for Three-Dimensional Visualization of Structures in the Cell (ed Frank, J.) 113–161 (Springer New York, New York, NY, 2006).

24. Frangakis, A. S. It’s noisy out there! A review of denoising techniques in cryo-electron tomography. Journal of Structural Biology 213, 107804 (Dec. 2021).

25. Cardone, G., Grünewald, K. & Steven, A. C. A resolution criterion for electron tomography based on cross-validation. Journal of Structural Biology 151, 117–129 (Aug. 2005).

26. Bhamre, T., Zhang, T. & Singer, A. Denoising and covariance estimation of single particle cryo-EM images. Journal of Structural Biology 195, 72–81 (July 2016).

27. Sindelar, C. V. & Grigorieff, N. An adaptation of the Wiener filter suitable for analyzing images of isolated single particles. Journal of Structural Biology 176, 60–74 (Oct. 2011).

28. Weiss, A. & Nadler, B. “Self-Wiener” filtering: Data-driven deconvolution of deterministic signals. IEEE Transactions on Signal Processing 70, 468–481 (2022).

29. Schaffer, M. et al. A cryo-FIB lift-out technique enables molecular-resolution cryo-ET within native Caenorhabditis elegans tissue. Nature Methods 16, 757–762 (Aug. 2019).

30. Buchholz, T.-O., Jordan, M., Pigino, G. & Jug, F. Cryo-CARE: Content-Aware Image Restoration for Cryo-Transmission Electron Microscopy Data in 2019 IEEE 16th International Symposium on Biomedical Imaging (ISBI 2019) (IEEE, Apr. 2019), 502–506.

31. Dietrich, H. M. et al. Membrane-anchored HDCR nanowires drive hydrogen-powered CO2 fixation. Nature 607, 823–830 (July 2022).

32. Moscovich, A., Halevi, A., Andén, J. & Singer, A. Cryo-EM reconstruction of continuous heterogeneity by Laplacian spectral volumes. Inverse Problems 36, 024003 (Feb. 2020).

33. Donnat, C., Levy, A., Poitevin, F., Zhong, E. D. & Miolane, N. Deep generative modeling for volume reconstruction in cryo-electron microscopy. Journal of Structural Biology 214, 107920 (Dec. 2022).

34. Toader, B., Sigworth, F. J. & Lederman, R. R. Methods for cryo-EM single particle reconstruction of macromolecules having continuous heterogeneity. Journal of Molecular Biology, 168020 (Feb. 2023).

35. Liu, Y.-T. et al. Isotropic reconstruction for electron tomography with deep learning. Nature Communications 13, 6482 (Oct. 2022).

36. Bepler, T., Kelley, K., Noble, A. J. & Berger, B. Topaz-Denoise: general deep denoising models for cryoEM and cryoET. Nature Communications 11, 5208 (Oct. 2020).

37. Li, H. et al. Noise-Transfer2Clean: denoising cryo-EM images based on noise modeling and transfer. Bioinformatics 38, 2022–2029 (Mar. 2022).

38. Lawson, C. L. et al. EMDataBank unified data resource for 3DEM. Nucleic Acids Research 44, D396–D403 (Jan. 2016).

39. Wright, G. et al. ComputationalCryoEM/ASPIRE-Python: v0.12.0 10.5281/zenodo.5657281 version v0.12.0. Sept. 2023.

40. Oppenheim, A. V., Willsky, A. S., Nawab, S. H., Hernández, G. M., et al. Signals & systems (Pearson Educación, 1997).

